# Membrane permeability selection drove the stereochemistry of life

**DOI:** 10.1101/2024.04.23.590732

**Authors:** Olivia Goode, Urszula Łapińska, Georgina Glover, David S. Milner, Alyson E. Santoro, Stefano Pagliara, Thomas A. Richards

## Abstract

Early in the evolution of life a proto-metabolic network was encapsulated within a membrane compartment. The permeability characteristics of the membrane determined several key functions of this network by determining which compounds could enter the compartment and which compounds could not. One key feature of known life is the utilisation of right-handed D- ribose and deoxyribose sugars and left-handed L- amino acid stereochemical isomers (enantiomers), however, it is not clear why life adopted this specific chirality. We previously demonstrated that an archaeal and an intermediate membrane mimic, bearing a mixture of bacterial and archaeal lipid characteristics (a ‘hybrid’ membrane), display increased permeability compared to bacterial-like membranes. Here, we investigate if these membranes can drive stereochemical selection on pentose sugars, hexose sugars and amino acids. Using permeability assays of homogenous unilamellar vesicles, we demonstrate that both membranes select for D- ribose and deoxyribose sugars while the hybrid membrane uniquely selects for a reduced alphabet of L- facing amino acids. This repertoire includes alanine, the plausible first L- amino acid utilised. We conclude such compartments could provide stereochemical compound selection thereby demonstrating a solution to the chirality problem during the evolution of life.

## Introduction

Life on Earth is defined by a curious and universal stereochemical asymmetry [1, 2]. Specifically, the pentose sugars utilised for DNA and RNA (deoxyribose and ribose) possess three stereocenters imposing a D- (right-handed) stereochemistry. In contrast, 19 of the 20 universal proteinogenic amino acids used possess one or two stereocenters imposing L- (left-handed) stereochemistry on polypeptides. A number of abiotic processes have been proposed for generation of homochiral states (e.g., [1–4]). Yet importantly, autocatalytic polymerisation of mononucleotides proceeds with efficacy in mixtures with high relative ratios of one enantiomer [5]. Similar catalytic processes have been demonstrated for amino acids which polymerise stereo-selectively in abiotic conditions [3, 6]. This demonstrates Wald’s conjecture [1], which set out how the polymerisation characteristics of ribonucleotides and amino acids would lead life to preferentially adopt a single stereochemical form. While this explains why life might use homochiral states, it does not explain the conditions under which specific homochiral mixes arose in a biologically proximate system allowing this bias to become hardwired into the central metabolism of life.

Ideas about the origin of life have been shaped by the demonstration that abiotic chemical conditions can generate many compounds utilised by living systems. Compounds generated by these abiotic processes are equal mixes of both enantiomers (racemic). Experimentally simulated prebiotic conditions have been shown to form ribose through the formose reaction [7]. The distribution of variant glycolysis pathways across the tree of life [8, 9] also suggests that glycolysis was an ancestral feature of the last universal common ancestor (LUCA) [10, 11]. Consistent with the phylogenetic age of these pathways, experimental chemical simulations of the early Earth Ocean led to the formation of glycolysis intermediates [12], suggesting pathway intermediates were available for catabolism and as precursors of ribonucleotides and phospholipids. Additional work has suggested that gluconeogenesis preceded glycolysis [13] and, therefore, glucose, and possibly other hexose sugars, were available for early life forms.

Similarly, the Miller experiments [14, 15] demonstrate that a subset of amino acids will form in simulated early Earth conditions. Importantly, these experiments demonstrate glycine and alanine are generated at much higher relative concentration than any other proteinogenic amino acids. Similar amino acid composition has been detected in meteorite-derived materials [16]. The meteorite data provide important ‘outside the lab’ demonstrations that chemical compounds relevant for proto-metabolic function can form in abiotic conditions.

A systematic review of 40 different criteria has been used to reconstruct the relative temporal order of amino acid utilisation as life evolved, providing a consensus that glycine and alanine were the first amino acids utilised [17]. Glycine does not possess any stereocenters, while alanine has one. As such, the use of L- alanine over D- alanine putatively imposed L- stereochemistry on all future biogenic proteins. However, the amino acids generated from the Miller experiments and detected in meteorite samples are racemic, possessing equal shares of both enantiomers [14, 16], a mixture of compounds that would restrict the evolution of early metabolic functions [1, 3, 6] without some source of enantiomer selection.

J. B. S. Haldane discussed how life could have emerged ‘enclosed in an oily film’ [18], a scenario which would have separated early biological systems from abiotic mixtures present in the surrounding environment. Independent of whether life started as a heterotrophic form [19, 20] (i.e., the ‘Oparin-Haldane’ theory [21]) or an autotrophic form [12, 22–25], a critical step in the history of life involved the encapsulation of metabolic functions within a membrane [26, 27]. Acquisition of a membrane bestowed a proto-cellular form with the capacity for metabolite accumulation [28], generation of chemical gradients [29], and co-association of interacting systems necessary for co-function [27]. Compartmentalisation also generated divisible units, allowing variation in biochemical functions including different reaction kinetics, upon which Darwinian selection could act [30, 31]. Cell membranes have been known to be semi-permeable since the late 18^th^ century [32, 33], forming a barrier that can selectively allow some compounds to pass through while excluding others. The permeability of proto-cellular membranes must therefore have determined the degree to which internal membrane space is a chemically distinct environment and therefore the evolution of early metabolic functions [28, 34, 35].

Fatty acid membrane compartments of various compounds are useful model systems [36] for understanding protocell evolution. For example, they have been shown to assemble from meteorite extracts [37] and can be generated from clays in a process which can encapsulate RNA templates [38]. Such membranes are also permeable to chemical precursors necessary for core cellular functions [19, 28, 35, 39] and can ‘grow’ in a process driven by vesicle content [30, 40] including internalised RNA replication [19]. Nonetheless, there are chemical limitations to fatty acid membranes as proto-cellular systems; chiefly, such vesicles would not maintain integrity in environments with high concentrations of divalent cations (e.g., seawater) [31]. Similarly, they could not host the high concentrations of Mg^2^+ thought necessary for ribozyme-catalysed polymerase function, which has been suggested to act as an early replicator in the RNA world hypothesis [19, 41, 42]. These observations suggest that a phase of evolution using an alternative membrane chemistry was necessary.

The membrane of all extant cellular life is built from variant phospholipids [43] Experiments have demonstrated that vesicles of eukaryotic membrane analogs (POPC and DPPC) are permeable to a range of pentose sugars including ribose [35], while vesicles of bacterial membrane chemical analogs (EPC and DMPC) are permeable to amino acids [34]. Phospholipid vesicles can encapsulate macromolecules upon dehydration-hydration cycles [44], making them important models for understanding how proto-cellular compartments could internalise metabolic compounds, templates or catalysts. Furthermore, phospholipids can form in abiotic conditions [45], can be used to encapsulate RNA polymerisation [46], can perform budding and division in the presence of *ad hoc* precursors [47], and can withstand ion concentration gradients [48].

A range of phospholipid forms are used by microbes, often in heterogenous and varying mixes. Despite this diversity, there are two fundamental core membrane phospholipid chemistries present in cellular life [43]: bacteria and eukaryotes possess membranes predominantly composed of fatty acid chains bonded to a glycerol-3-phosphate (G3P) backbone via ester bonds; a phospholipid with a right-handed stereochemistry. In contrast, most archaeal membranes are predominantly constructed from diether lipids with isoprenoid chains containing methyl branches bonded to a glycerol-1-phosphate (G1P) backbone via ether bonds; a phospholipid with a left-handed stereochemistry [43, 48, 49]. This pattern demonstrates that deep within the tree of life two different lipid forms were adopted on distinct branches [43]. It is not clear if this transition was from: i) one form to the other, ii) an intermediate form (for example, a phospholipid with a combination of bacterial and archaeal characteristic i.e., a ‘hybrid’ membrane), iii) the fatty acid membranes similar to the types discussed above, iv) the common ancestor of Archaea and Bacteria possessed a mixed heterochiral membrane, or v) an intermediate that possessed no membrane; with the two membrane types adopted later and independently [24, 50, 51]. There is currently limited consensus on the placement of the last universal common ancestor (LUCA) which would enable us to polarise the transition in membrane chemistry. Some analyses suggests that the root of the tree of life may reside between the Bacteria and the Archaea [52–57], whilst alternative analyses suggest that the root lies within the Bacteria [58–60]. The placement of the root determines the minimum number and nature of transitions in membrane chemistry during the evolution of the prokaryotic domains, however, neither rooting scenario excludes presence of hybrid membrane variants deep within either the Bacteria or the Archaea or at their common ancestor. Such hybrid forms may have had a critical impact on permeability characteristics during the early evolution of cellular forms.

Using unilamellar vesicle microfluidic permeability assays, we have previously shown that archaea-like G1PC phospholipid membranes demonstrate higher permeability to core metabolites than bacterial-like membranes. These core metabolites include: nucleobases (that possess no stereochemistry), ribose, deoxyribose and six proteinogenic amino acids [61]. These experiments also included comparisons with a plausible intermediate ‘hybrid’ phospholipid membrane consisting of diether lipids with isoprenoid chains containing methyl branches (archaeal traits) bonded to a glycerol-3-phosphate (G3P) with right-handed stereochemistry (bacterial traits). Experiments using this hybrid membrane demonstrated increased permeability compared to both bacterial and archaeal membranes for deoxyribose, ribose and glycine. Alternative hybrid membrane forms with different combinations of chemical characteristics showed reduced permeability in comparison to both the hybrid and archaeal membrane. We therefore suggest that the permeability characteristics of archaeal-like membranes might have driven the early evolution of proto-cellular metabolism [61].

Historically, few experiments have compared L- and D- enantiomer permeability through membranes. For example, Nagano in 1902 [62, 63] compared the absorption rate of a range of sugars across the intestinal membrane of dogs, an experiment that included comparisons of D- hexose to L- pentose sugars. More recently, Sacerdote and Szostak compared absorption of L- and D- xylose through four homogeneous vesicle types, including three different fatty acids and one phospholipid (POPC, a compound similar to eukaryotic membranes) [35]. This experiment demonstrated no difference in the enantiomer permeability of xylose. Here, we expand such comparisons to a range of compounds useful for proto-metabolism and focus on archaeal and hybrid membrane mimics. We focus on these two membrane mimics because our previous experiments show they display the highest permeability [61]. The aim of this work is to test the hypothesis that membrane chemistry imposes permeability selection on enantiomers. We compare the enantiomer permeability of four pentose sugars, two hexose sugars and twelve amino acids, demonstrating strong selection for D- ribose and D- deoxyribose in both membrane types. In contrast only the hybrid membrane shows permeability selection for a handful of L- amino acids including strong selection for L- alanine. We therefore suggest that early utilisation of a ‘hybrid’ membrane chemistry –a possible precursor form to either or both bacterial or archaeal membrane chemistry– would have allowed enrichment of specific enantiomers and therefore hardwired a proto-cellular form to utilise L- form amino acids and D- form ribonucleotides.

## Results

### Pentose sugar permeability shows selection for D- ribose and D- deoxyribose

To explore the possibility that an archaeal or hybrid membrane imposes permeability selection on different enantiomers we compared permeability of four pentose sugars: deoxyribose, ribose, xylose and arabinose (Figure 1). Both membranes demonstrate permeability to ribose, xylose and arabinose and, importantly, show increased permeability to D- ribose compared to L- ribose (P < 0.01). Stereochemical selection was not evident for xylose or arabinose.

**Figure 1.** Membrane permeability to pentose sugars shows bias for D- ribose and D- deoxyribose. Temporal dependence of average carboxyfluorescein (CF) fluorescence in the archaea-like G1PC phospholipid membrane (G1PC, filled symbols) and in the ‘hybrid’ phospholipid membrane (G3PC, open symbols) during the exposure to 1 mM of variant substrates delivered to the microfluidic coves at constant rate. Mean (symbols) and standard deviation (error bars) were calculated from at least 10 single-vesicle measurements across three independent experiments. Lines connecting points are provided as a guide. N is the number of single vesicles investigated for each substrate exposure and each type of vesicle. N varies across different substrate experiments investigated due to technical constraints. However, care has been taken to obtain the same N for each substrate experiment across the two different enantiomer treatments in order to ensure reliable statistical comparisons. Such comparisons have been carried out via Welch’s t-tests between the distributions of CF fluorescence values at t = 3 min for each enantiomer pair and are shown with corresponding violin plots next to each time-course graph (C, F, I, L). ****: p-value < 0.0001, **: p-value < 0.01. Numerical values of CF fluorescence in individual vesicles for each lipid type during the delivery of each substrate are provided in S1 File and S2 File.

As discussed above, permeability for D- ribose has been reported for both fatty acid membranes and some eukaryotic-like phospholipids, but L- and D- rates were not compared as part of these experiments. However, those experiments did include a comparison of permeability for L- and D- xylose [35], demonstrating: i) reduced permeability for both enantiomers of xylose compared to D- ribose, and ii) no difference in permeability between the two xylose enantiomers through POPC eukaryotic-like phospholipids [35] similar to our results for xylose in both the membranes tested here (Figure 1). Our experiments therefore also produce data consistent with the idea that ribose was favoured over xylose or arabinose by membrane selection (Figure 1, & S1).

L- deoxyribose showed very low average permeability through either membrane type but, in contrast, D- deoxyribose showed significantly increased permeability through both membrane types (P < 0.01 and P < 0.0001 for the archaeal and hybrid membrane, respectively). These results demonstrate that both the archaeal and hybrid membrane can drive stereochemical selection generating a ‘passive sorting mechanism’ [35] for D- ribose and D- deoxyribose. Equivalent comparisons of bacterial membrane mimics do not show stereochemical selection (Figure S1). The patterns of stereochemical selection observed through the archaeal and hybrid membranes are consistent with the utilisation of these compounds as the chemical building blocks for DNA and RNA, the molecules central to the hereditary, transcription and translational systems of all life [64, 65].

### Hexose sugar permeability through archaeal and hybrid membranes

As discussed above, another common function that likely emerged early in the evolution of life is the glycolysis pathway [11, 12, 49]. This pathway provides chemical precursors for both RNA and phospholipid compounds, suggesting a varied role in the evolution of life [50]. Although there is considerable differentiation in the glycolysis pathways across the tree of life, and in many cases enzymes of the pathway are not encoded by homologous genes [8, 9], both archaea and bacteria can use glucose and fructose as a primary carbon source [66–68].

To understand how early cellular forms could make use of abiotically derived glucose or fructose [12, 13] (both with four stereocenters), we performed permeability selection assays through the archaeal and hybrid membranes. These data show both hexoses can permeate through these membrane forms; but a minor (P < 0.01), selection for D- fructose over L- fructose enantiomers was observed through the archaeal membrane (Figure 2). We found no selection for L- or D- fructose enantiomers through a bacterial membrane (Figure S1). These data suggest utilisation of the archaeal membrane could have generated a bias for a D- fructose precursor providing a D- source metabolite bias for glycolytic metabolism.

**Figure 2.** Hexose sugar permeability through archaeal and hybrid membranes. Temporal dependence of average carboxyfluorescein (CF) fluorescence in the archaea-like G1PC phospholipid membrane (G1PC, filled symbols) and in the ‘hybrid’ phospholipid membrane (G3PC, open symbols) during the exposure to 1 mM of variant substrates delivered to the microfluidic coves. Mean (symbols) and standard deviation (error bars) were calculated from at least 10 single-vesicle measurements across three independent experiments. Data are processed and illustrated as described in Figure 1 legend. N is the number of single vesicles investigated for each substrate exposure and each type of vesicles. **: p-value < 0.01. Numerical values of CF fluorescence in individual vesicles for each lipid type during the delivery of each substrate are provided in S1 File and S2 File.

### Amino acid permeability through archaeal and hybrid membranes

We previously demonstrated that archaeal membranes have a higher permeability to glycine and alanine compared to bacterial membranes [61]. To explore if the archaeal and the hybrid membranes impose selection on alanine, we compared permeability of L- and D- alanine enantiomers. We found no differential selection through the archaeal membrane, but strong selection for the L- alanine enantiomer in comparison to the effective exclusion of D- alanine in the hybrid membrane (P < 0.0001, Figure 3). We also note that we had previously demonstrated that L- alanine does not permeate through bacterial membranes [61]. This result demonstrates a passive L- alanine sorting mechanism consistent with the amino acid stereochemical asymmetry early in the evolution of amino acid proteinogenic utilisation, but only in the hybrid membrane and not the archaeal membrane mimic tested.

**Figure 3.** Archaeal and hybrid membrane permeability to non-polar amino acids. Temporal dependence of average carboxyfluorescein (CF) fluorescence in the archaea-like G1PC phospholipid membrane (G1PC, filled symbols) and in the ‘hybrid’ phospholipid membrane (G3PC, open symbols) during the exposure to 1 mM of variant substrates delivered to the microfluidic coves. Data are processed and illustrated as described in Figure 1 legend. Mean (symbols) and standard deviation (error bars) were calculated from at least 10 single-vesicle measurements across three independent experiments. N is the number of single vesicles investigated for each substrate exposure and each type of vesicles. ****: p-value < 0.0001, **: p-value < 0.01. Numerical values of CF fluorescence in individual vesicles for each lipid type during the delivery of each substrate are provided in S1 File and S2 File.

To further explore stereochemical permeability selection, we performed permeability assays on 11 additional proteinogenic amino acids. These 11 amino acids included seven amino acids (V, D, E, S, L, T, I) that were shown to be produced from the Miller experiments but at much lower molar concentrations than alanine or glycine [14, 15]. The amino acids analysed include compounds with two stereocenters (T, I) and a range of differing physical-chemical properties including: three non-polar (V, L, I, Figure 3), five polar (S, T, N, C, Q, Figure 4), one positively and two negatively charged amino acids (R, D, E, respectively, Figure 5). These assays confirmed that all 14 amino acids tested (when including glycine and tryptophan previously analysed [61]) permeated through the archaeal membrane (Figures 3-5). Furthermore, focusing on amino acids where enantiomers were compared: I, S, C, R and D amino acids demonstrated enantiomer selection through the archaeal membrane. However, these results were mixed, with only arginine (R) showing L- selection while the other four (D, C, S, I) showed D- selection (Figures 3-5). This pattern is not consistent with how life evolved to use amino acid enantiomers.

**Figure 4.** Polar amino acid permeability through archaeal and hybrid membranes. Temporal dependence of average carboxyfluorescein (CF) fluorescence in the archaea-like G1PC phospholipid membrane (G1PC, filled symbols) and in the ‘hybrid’ phospholipid membrane (G3PC, open symbols) during the exposure to 1 mM of variant substrates delivered to the microfluidic coves. Data are processed and illustrated as described in Figure 1 legend. Mean (symbols) and standard deviation (error bars) were calculated from at least 10 single-vesicle measurements across three independent experiments. N is the number of single vesicles investigated for each substrate exposure and each type of vesicles. ****: p-value < 0.0001, *: p-value < 0.05. Numerical values of CF fluorescence in individual vesicles for each lipid type during the delivery of each substrate are provided in S1 File and S2 File.

**Figure 5.** Charged amino acid permeability through archaeal and hybrid membranes. Temporal dependence of average carboxyfluorescein (CF) fluorescence in the archaea-like G1PC phospholipid membrane (G1PC, filled symbols) and in the ‘hybrid’ phospholipid membrane (G3PC, open symbols) during the exposure to 1 mM of variant substrates delivered to the microfluidic coves. Data are processed and illustrated as described in Figure 1 legend. Mean (symbols) and standard deviation (error bars) were calculated from at least 10 single-vesicle measurements across three independent experiments. N is the number of single vesicles investigated for each substrate exposure and each type of vesicles. ***: p-value < 0.001, *: p-value < 0.05. Numerical values of CF fluorescence in individual vesicles for each lipid type during the delivery of each substrate are provided in S1 File and S2 File.

In contrast, stereochemical permeability assays through the hybrid membrane (a phospholipid with the opposite D- stereochemistry than the archaeal membrane [43]) demonstrated fewer amino acids could permeate this membrane. Specifically, C, E, S, T and V showed no permeability in contrast to the archaeal membrane (Figures 3-5). Consistent with the alanine results discussed above, the hybrid membrane showed L- enantiomer selection for L, Q, R, and D amino acids (Figures 3-5), demonstrating how this membrane can generate passive selection for a reduced alphabet of L- amino acids. These results show that specific chemical characteristics of the membrane phospholipid can determine membrane permeability function. We also note, as the hybrid membrane effectively excluded multiple amino acid enantiomers, these experiments confirm that the amino acid selectivity represents true differences in membrane permeability function (see also Figure S3 for further controls).

To test for enantiomer selection through the bacterial membrane we compared L- and D- permeability of R, C, and L amino acids through a bacterial mimic. These amino acids were chosen as they showed very different enantiomer permeability through archaeal and hybrid membranes (Figure 3-5). These experiments demonstrate no evidence of enantiomer selection through the bacterial membrane (Figure S2).

## Discussion

Using controlled unilamellar vesicle experiments, we demonstrate that a model archaeal membrane and one hybrid membrane form drive stereochemical permeability selection (summarised in Table 1). These data suggest an ancient phase of evolution based on either of these two-membrane chassis could have driven selection for the stereochemical characteristics of proto-metabolism. Specifically, an evolutionary bottleneck dependent on either membrane chemistry tested would have generated a high level of selection for D-ribose and D- deoxyribose, hard-wiring the use of right-handed ribonucleotides. Strikingly, a cell-type with a hybrid membrane -a possible precursor of both archaeal and bacterial membrane types-could have hardwired both the use of D- ribonucleotide oligonucleotides and L- form amino acids, although initially with a reduced alphabet of amino acids. Given current data, we advocate for a phase of evolution prior to, or at the last common ancestor, which was a hybrid membrane consisting of diether lipids with isoprenoid chains containing methyl branches (archaeal traits) bonded to a glycerol-3-phosphate (G3P) with right-handed stereochemistry (bacterial traits). We note that this scenario only represents one phase of membrane evolution in a long history of possible membrane transitions between the first ‘life’ form and the last common ancestor. As such this scenario is compatible with many previously suggested ancestries for the prokaryotic domains (e.g., [24, 50, 51]).

**Table 1.**
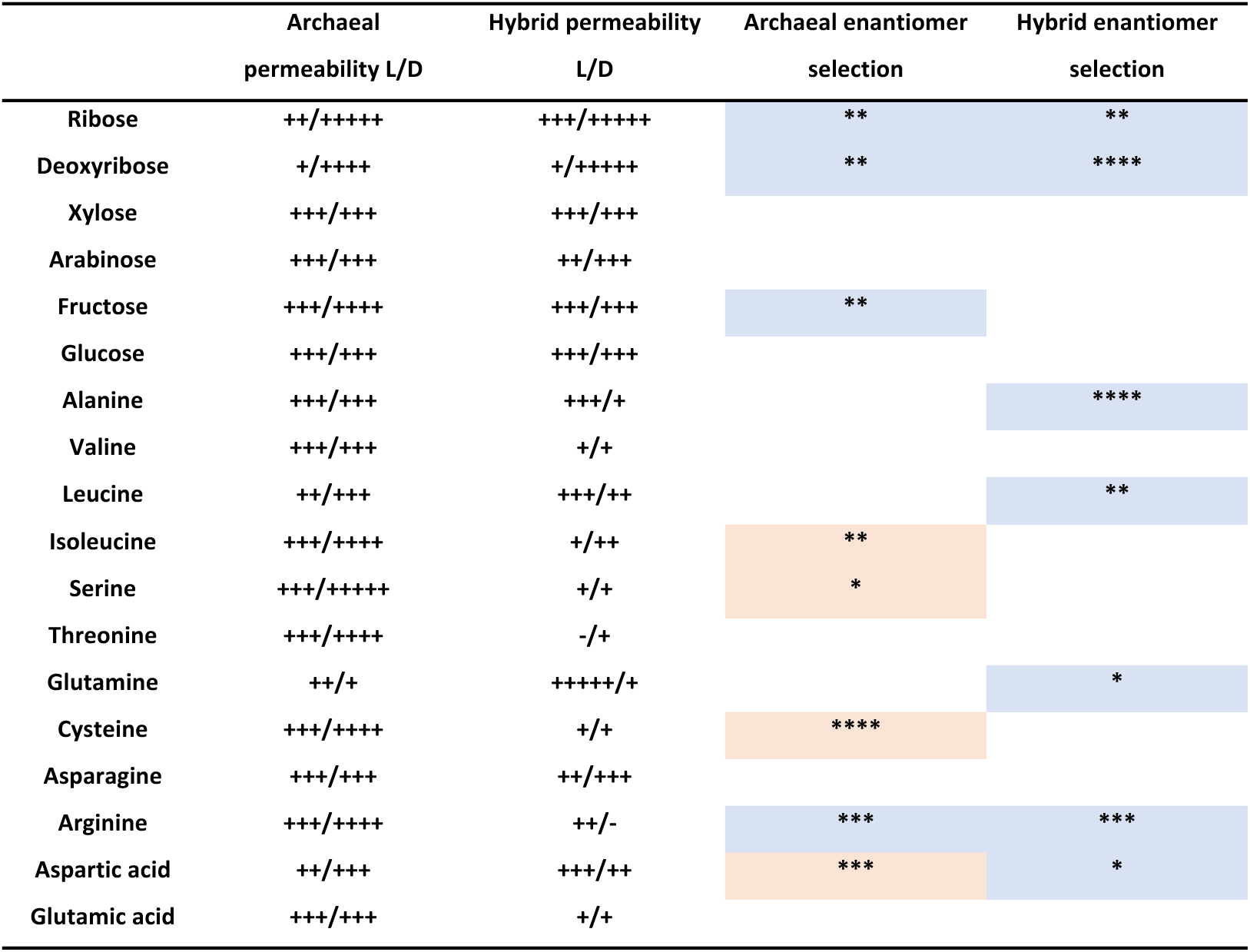
Summary of permeability and enantiomer selection data for the eighteen substrates tested against the two membranes. Carboxyfluorescein (CF) fluorescence after 3 minutes of exposure to each substrate at a concentration of 1 mM is summarised as follows: <0 = -; 0-5 a.u. = +, 5-10 a.u. = ++, 10-15 a.u. = +++, 15-20 a.u. = ++++, >20 a.u. = +++++. Statistical support for different enantiomer selection is indicated: ****: p-value < 0.0001, ***: p-value < 0.001, **: p-value < 0.01,*: p-value < 0.05. Enantiomer selection consistent with the stereochemical bias of life is coloured blue. Enantiomer selection that contradicts the stereochemical bias of life is coloured orange.

Different phylogenomic analyses have suggested that the universal common ancestor lies between the Archaea and the Bacteria [52–57], or potentially within the Bacteria [58–60]. Our results do not directly inform this debate, as multiple transitions in membrane chemistry are possible; indeed, the transition through the hybrid membrane we advocate could have occurred before the derivation of either the Archaea or the Bacteria. It is also important to mention that a pure form membrane, as used here, is implausible as an early proto-cellular chassis, especially if they were derived from an abiotic source [45] and occurred prior to the emergence of a dedicated metabolic network for the generation of membrane forming compounds. It is not clear, at this stage, if mixed membranes encompassing the types of phospholipids used here are viable, although experiments with *E. coli* suggest glycerol-1-phosphate and glycerol-3-phosphate lipids can form a heterochiral membrane [69]. Furthermore, if it is possible to form mixed phospholipid unilamellar vesicles, would such membranes generate similar patterns of permeability selection identified here. Future works should focus on assessing the stability and permeability function of mixed membranes.

The results reported here also demonstrate that additional bottlenecks through an archaeal type membrane could have hard-wired bias for D-hexose sugars for metabolic utilisation such as glycolysis, as demonstrated by our result of increased archaeal membrane permeability to D-fructose (Figure 2). However, this was not a strong result, and the possibility of two bottlenecks with differing membrane chemistries leaves some confusion regarding the relative inferred order of membrane utilisation relative to the tree of life. We also note that glycolytic catabolism of hexose sugars does not impose similar constraints on enantiomer utilisation as would the emergence of the anabolic polymerisation of ribonucleic acids and amino acid polypeptides [1]. This is because hexose compounds are primarily broken down for metabolic benefit and not polymerised for storage or translation of information in the form of DNA, RNA and amino acid polypeptides. Differential utilisation, i.e., polymerisation in the case of amino acids and ribonucleotides vs catabolic metabolism in the case of glycolysis of hexoses, therefore suggests that the stereochemical permeability selection on precursors of glycolysis is not as an important a factor as precursors of the compounds utilised in the central dogma.

We acknowledge that wider analysis of membrane types, including bacterial-mimics may reveal additional permeability selection traits informative for understanding proto-cellular membrane evolution. We have partly investigated this possibility in a bacterial membrane mimic for three pentose sugars, one hexose sugar and three amino acids (Figure S1-2). These were chosen as they showed a wide variation of permeability characteristics in the archaeal and the hybrid membranes tested (Table 1). In all these tests the bacterial membrane mimic demonstrated no stereochemical selection, an outcome that could be a function of their lower permeability identified on our experimental platform [61].

Here we have discussed published examples of chemical polymerisation of nucleotides and amino acids in abiotic conditions [1, 3, 5, 6]. In these examples both types of polymerisation are limited when conducted in racemic mixes of substrate due to enantiomer compatibility and the reducing availability of compatible enantiomers as polymerisation progresses in a closed pool of chemical precursors. The membrane permeability selection demonstrated here would solve this problem by providing a compartment with specific enantiomer enrichment, allowing polymerisation to progress without inhibition generated by racemic mixes of the constituent subunits. Such a system would have multiple advantages in competition over protocells without enantiomer selection. Specifically, both permeability and polymerisation selection would act synergistically to generate competitive advantage and therefore Darwinian selection [30, 31]. Understanding how life began is a jigsaw puzzle [70]; the utilisation of a hybrid phospholipid membrane to derive stereochemical selection for deoxyribose, ribose and a reduced alphabet of amino acids (Table 1) is a potential piece of that puzzle. These observations are valuable information towards explaining the chirality problem as part of the origin of life, however we acknowledge that these observations do need testing with alternative methods.

## Methods

All methods are adaptations of those reported in Łapińska et al. 2023 [61].

### Preparation of materials

All chemicals were purchased from Merck. All Lipids were purchased from Avanti Polar Lipids [i.e. 1,2-di-O-phytanyl-sn-glycero-1-phosphocholine, 1,2-di-O-phytanyl-sn-glycero-3-phosphocholine, 1,2-dioleoyl-sn-glycero-3-phosphoethanolamine, 1,2-dioleoyl-sn-glycero-3-phospho-(1’-rac-glycerol) and 1,3-bis(sn 3’-phosphatidyl)-sn-glycerol)]. Indium tin oxide (ITO)-coated glass slides were purchased from VisionTek Systems. The fluorescent probe 5(6)-carboxyfluorescein (CF), (MW=376 g/mol), was dissolved in absolute ethanol at a stock concentration of 10 mg/mL. In order to perform all permeability experiments at physiological pH (7.4), the washing buffer was prepared by dissolving sucrose (MW = 342 g/mol) in 5 mM 4-(2-hydroxyethyl)-1-piperazineethanesul-fonic acid (HEPES) at pH=7.4 at a final sucrose concentration of 195 mM. All metabolites were dissolved in the washing buffer to a final concentration of 1 mM.

### Preparation of synthetic vesicles

Giant unilamellar vesicles were electroformed using a Vesicle Prep Pro (Nanion) [71]. Three different lipid stocks; archaeal lipid mimic (G1PC) (2,3-di-O-phytanyl-sn-glycero-1-phosphocholine), hybrid lipid mimic (G3PC) (1,2-di-O-phytanyl-sn-glycero-3-phosphocholine) and bacterial lipid mimic mixed of three lipids: 1,2-dioleoyl-sn-glycero-3-phosphoethanolamine, 1,2-dioleoyl-sn-glycero-3-phospho-(1’-rac-glycerol) and 1,3-bis(sn-3’-phosphatidyl)-sn-glycerol were prepared to a final concentration of 10 mM dissolved in chloroform (Table S1.). Next, 10 µL of each lipid solution was spread evenly onto the conductive side of an ITO-glass slide and vacuum desiccated to dry the solution. Meanwhile, 495 µL of washing buffer was degassed and mixed with 5 µL of 10 mg/ml of 5(6)-CF solution to a final concentration of 0.266 mM. The lipid coated ITO-glass slide was loaded into the Nanion and a greased rubber O-ring was placed over the dried layer. The area inside the O-ring was filled with 300 µL of 0.266 mM CF solution and another ITO-glass slide conductive side down was placed over, avoiding the formation of bubbles. Different electroformation protocols were implemented depending on the lipid used (see Table S2.). The final suspension contains CF fluorescent vesicles and free CF molecules, which were stored at 4°C for 2 days maximum.

### Fabrication of the microfluidic device

The microfluidic device is described in detail in Łapińska, 2023 [61]. In short, the device contains two inlets connecting to a main chamber split into 4 channels, these contain hydrodynamic traps (also known as coves) with the chambers finally leading to a single outlet. Using a mould created by multilevel photolithography [72], polydimethylsiloxane (PDMS) replicas of the device were produced. In summary, a 10:1 (base:curing agent) PDMS mixture (SYLGARD 184 Silicone Elastomer Kit, Dow) was poured over the mould and degassed for 30 mins, this was cured for 2 hr at 70°C in an oven. The PDMS was peeled from the mould and a 1.5 mm biopsy punch was used to create fluidic access. To irreversibly seal the PDMS replica to a glass slide, oxygen plasma treatment was used (10 sec exposure to 30W plasma power, Plasma etcher, Diener Electronic GmbH).

### Microfluidic permeability assay

Washing buffer was injected into the buffer inlet until the device was flooded completely and escaping the metabolite inlet and outlet. The metabolite inlet was then sealed with tape and 10 µL of the fluorescent vesicle suspension was pipetted into the washing inlet. The tape prevented fluorescent vesicle suspension entering the metabolite inlet. The chip was mounted onto an inverted epifluorescence microscope (Olympus IX73) equipped with a sCMOS camera (Zyla 4.2, Andor, used at an exposure time of 0.1s), a blue LED (CoolLED pE300white, used at 30% of its intensity) and a FITC filter. Prepared 1 mL syringes were filled with washing buffer, and the second with 1 mM metabolite of interest were connected to 23-gauge needles (Becton Dickinson) and Masterflex Transfer Tygon tubing with 0.5 mm inner and 1.5 mm outer diameter (Cole-Parmer Instrument). These were controlled by a Nemesys pump operated via the Qmix Elements software (Centoni). The tape was removed from the metabolite inlet and the tubing containing the metabolite of interest was inserted flowing at a rate of 0.5 µL/hr. Then, the tubing containing the washing buffer was inserted into the remaining inlet, flowing at a rate of 5 µL/hr. These flow rates were increased in a stepwise fashion at steps of 0.5 µL/hr up to 2 µL/hr for the metabolite solution and 5 µL/hr up to 25 µL/hr for the washing buffer. These were kept constant for 20 mins to remove any free CF molecules from the microfluidic device. After this time the metabolite solution was increased to 25 µL/hr and the washing buffer solution was reduced to 1 µL/hr. Brightfield and fluorescent imaging of an area containing 14 coves were acquired every 30 sec for 3 mins.

### Vesicle free carboxyfluorescein and metabolite interaction assays

To explore the possibility that compound stereochemical variation altered carboxyfluorescein function, we ran separate microplate reader experiments demonstrating that the stereochemistry of the metabolic compound does not have a differential effect on the fluorescence of carboxyfluorescein (Figure S3), the reporter used throughout these experiments. These data provide a further control for our permeability selection data reported above.

Washing buffer was used to dissolve CF to a concentration of 0.532 mM and 100 µL was added to wells of a 96-well plate (Costar 96, Fisher Scientific). Then, 100 µL of 1 mM metabolite solution was added to each well and the plate placed within a Clariostar Plus plate reader (BMG Labtech, UK) for fluorescence data acquisition over 30 mins (Excitation 483-14, Emission 596, with a FITC filter). Three independent experiments were carried out for each metabolite.

### Image and data analysis

Images for each membrane/metabolite combination tested, at every timepoint, were exported into ImageJ for quantitative comparative analysis. Only single unilamellar vesicles were retained for analysis and a circle was drawn around each vesicle. A second identical circle was drawn in an area 10 µm upstream of the vesicle, which was then used to produce background-subtracted values for intra-vesicle fluorescence. The initial intra-vesicle fluorescence value (t=0) was subtracted. The values were then corrected for the impact of both the delivery of the washing buffer solution and photobleaching on the intra-vesicle CF fluorescence signal (see methods detailed in [73]). The permeability of each membrane mimic was measured in three independent experiments. Statistical comparisons were carried out using the Welch’s t test.

## Acknowledgements

We thank Nick Irwin for informative discussions relating to this work.

## Supplementary Tables and Figure Legends

**Table S1.**
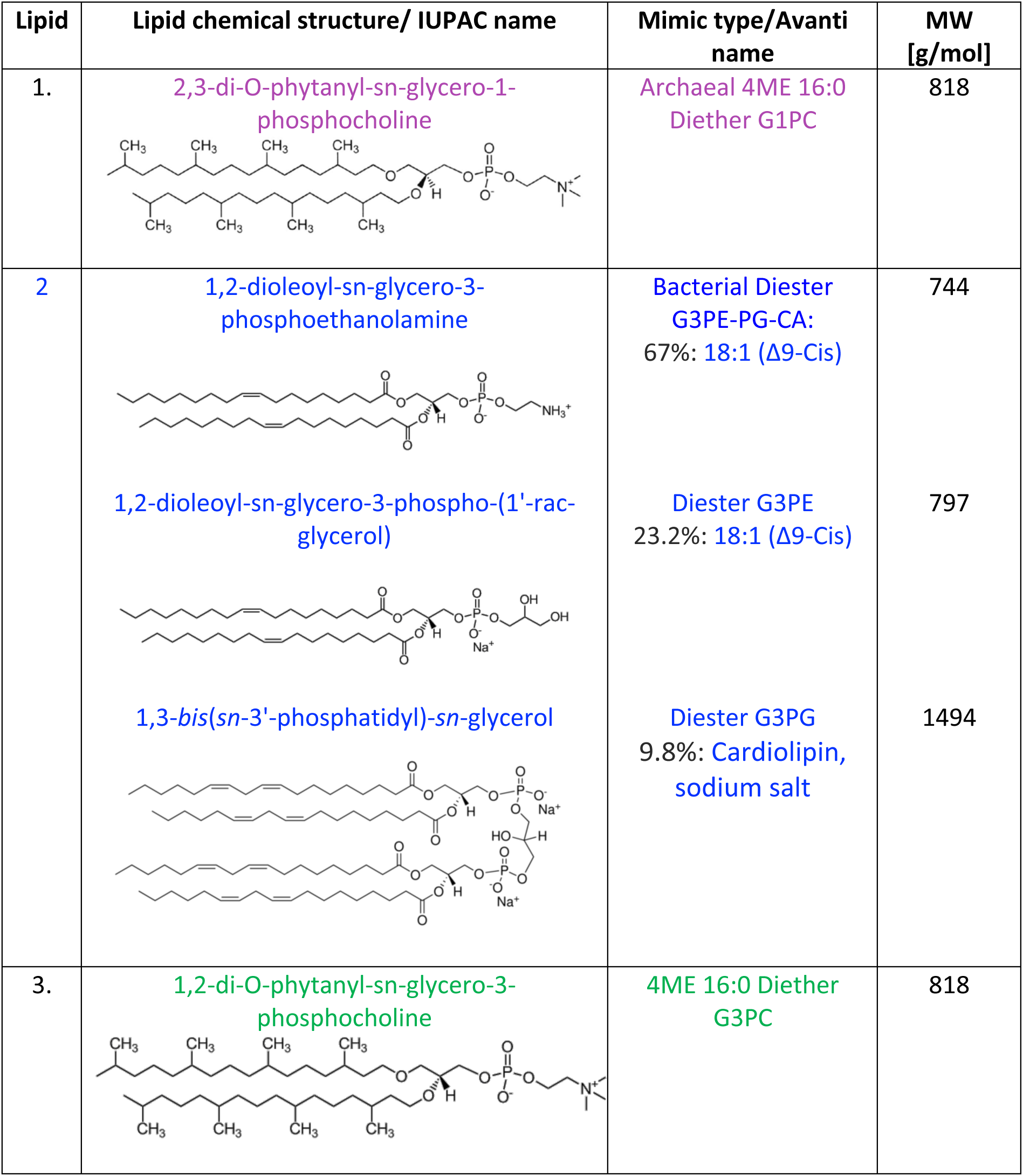
Details of membrane phospholipid compounds used for vesicle synthesis.

| Lipid | Lipid chemical structure/ IUPAC name | Mimic type/Avanti name | MW [g/mol] |
| --- | --- | --- | --- |
| 1. | 2,3-di-O-phytanyl-sn-glycero-1-phosphocholine | Archaeal 4ME 16:0 Diether G1PC | 818 |
| 2 | 1,2-dioleoyl-sn-glycero-3-phosphoethanolamine | Bacterial Diester G3PE-PG-CA: 67%: 18:1 (Δ9-Cis) | 744 |
|  | 1,2-dioleoyl-sn-glycero-3-phospho-(1'-rac-glycerol) | Diester G3PE 23.2%: 18:1 (Δ9-Cis) | 797 |
|  | 1,3-bis(sn-3'-phosphatidyl)-sn-glycerol | Diester G3PG 9.8%: Cardiolipin, sodium salt | 1494 |
| 3. | 1,2-di-O-phytanyl-sn-glycero-3-phosphocholine | 4ME 16:0 Diether G3PC | 818 |

**Table S2.** Parameters used for electroformation of vesicles.

| Lipid | Frequency<br>[Hz] | Amplitude<br>[V] | Temp.<br>[°C] | Time-rise<br>[min] | Time-main<br>[min] | Time-fall<br>[min] |
| --- | --- | --- | --- | --- | --- | --- |
| 1. & 3. | 5 | 3 | 37 | 5 | 120 | 5 |
| 2. | 10 | 1.6 | 37 | 5 | 160 | 5 |

**Figure S1. Sugar enantiomer permeability through bacterial membranes.** Temporal dependence of average carboxyfluorescein (CF) fluorescence in the bacteria-like phospholipid membrane during the exposure to 1 mM of variant substrates delivered to the microfluidic coves. Mean (symbols) and standard deviation (error bars) were calculated from at least 10 single-vesicle measurements across three independent experiments. Lines are guides for the eye. N is the number of single vesicles investigated for each substrate exposure. N varies across different substrate experiments investigated due to technical constraints. However, care has been taken to obtain the same N for each substrate experiment across the two different enantiomers treatments in order to ensure reliable statistical comparisons. Such comparisons have been carried out via Welch’s t tests between the distributions of CF fluorescence values at t = 3 min for each enantiomer and are shown with corresponding violin plots next to each time-course graph (B, D, F, H). ****: p-value < 0.0001, **: p-value < 0.01. Numerical values of CF fluorescence in individual vesicles during the delivery of each substrate are provided in S3 File.

**Figure S2. Amino acid enantiomer permeability through bacterial membranes.** Temporal dependence of average carboxyfluorescein (CF) fluorescence in the bacteria-like phospholipid membrane during the exposure to 1 mM of variant substrates delivered to the microfluidic coves. Mean (symbols) and standard deviation (error bars) were calculated from at least 10 single-vesicle measurements across three independent experiments. Lines are guides for the eye. N is the number of single vesicles investigated for each substrate exposure. N varies across different substrate experiments investigated due to technical constraints. However, care has been taken to obtain the same N for each substrate experiment across the two different enantiomers in order to ensure reliable statistical comparisons. Such comparisons have been carried out via Welch’s t tests between the distributions of CF fluorescence values at t = 3 min for each enantiomer and are shown with corresponding violin plots next to each time-course graph (B, D, F). ****: p-value < 0.0001, **: p-value < 0.01. Numerical values of CF fluorescence in individual vesicles during the delivery of each substrate are provided in S3 File.

**Figure S3. Metabolite stereochemistry does not have an impact on carboxyfluorescein fluorescence in solution.** Temporal dependence of (A) carboxyfluorescein (CF) fluorescence alone or in the presence of D- or L- enantiomers (red squares or blue circles, respectively) of (B-E) sugars, or (F-J) amino acids. Mean (symbols) and standard deviation (error bars) were calculated from three independent experiments performed in 96-well plates with fluorescence measured via a plate reader (note the reduced scale of the Y axis compared to the other plots reported). 100 µL of 0.532 mM CF was added to 100 µL of 1 mM metabolite; these were the concentrations used in the permeability experiments above. Numerical values of CF fluorescence in individual experiments in the presence of each metabolite are provided in S4 File.

